# An Integrated Computational Antigen Discovery Pipeline with Hierarchical Filtering for Emerging Viral Variants

**DOI:** 10.64898/2026.03.02.709067

**Authors:** Raj S. Roy, Jaemin Oh, A N M Nafiz Abeer, Maria I. Giraldo, Tesuro Ikegami, Scott C. Weaver, Nikolaos Vasilakis, Byung-Jun Yoon, Xiaoning Qian

## Abstract

Emerging and evolving viral diseases, such as SARS-CoV-2, continue to pose significant global health challenges, underscoring the urgent need for rapid and scalable antigen discovery pipelines. This work presents a computational pipeline that integrates diverse computational tools and machine learning models to accelerate the identification and optimization of antigen candidates. The pipeline employs efficient filtering and consensus-based strategies to highlight epitopes with high therapeutic potential. We demonstrate its utility by significantly narrowing the antigen search space for Rift Valley fever virus (RVFV) and Mayaro virus (MAYV), and by effectively identifying conserved neutralizing epitopes in SARS-CoV-2. Our proposed computational antigen pipeline offers a powerful framework for expediting the development of future vaccines and therapeutics in response to emerging pathogens.

## 1 Introduction

Emerging or circulating viral pathogens threaten human health. To address the challenges of widespread viral infections, both therapeutic and proactive strategies can be taken. Therapeutic strategies involve the design of drugs, antibodies, and nanobodies to effectively eliminate or neutralize viral agents in the human body. In contrast, proactive strategies include the design of vaccines. However, vaccine development remains a challenging task due to the significant cost and time required to validate its efficacy. Proactive strategies are generally more cost-efficient than therapeutic approaches, as they rely on the adaptive immune system to prevent infection and reduce the risk of reinfection.

Recent progress in computational resources and modeling techniques has facilitated the accurate prediction of protein structures and the design of novel proteins [1, 3, 11, 27, 31, 52]. In parallel with advancements in protein structure prediction and design, the development of machine learning (ML) algorithms and the extraction of highly informative features from large-scale biological datasets [4, 9] have enabled unprecedented accuracy in predicting diverse dynamics and properties relevant to immune responses. These include, for example, the identification of binding regions within antigen-antibody complexes [21, 26, 28, 32, 40], as well as the prediction of antigenicity and immunogenicity [14, 30], and the modeling of antibody escape evolution [19, 57]. One of the recent milestones of AI in therapeutics is the *de novo* snake antivenom, where RfD-iffusion played a key role in the design of the target binder [48]. A study reports that 80-90% of the AI-discovered molecules are successful in phase I of the clinical trial, and a 40% success rate in phase II [25]. This trend suggests that researchers are shifting towards AI-driven approaches as the traditional approaches are time-consuming and expensive [22].

However, there are few related works that combine existing computational tools to discover antigen in vaccine development. Rather, many works have focused on relatively narrow and specific tasks such as eptiope and property prediction. The few works related to the antigen pipeline are designed to combine multiple epitope prediction tools to design multi-epitope vaccines [10, 34, 36, 56]. Designs of this sort often require manual interventions, exhibit excessive dependency in individual tool and structure-based docking [12, 37, 55], which are computationally taxing and slow. Furthermore, such approaches frequently overlook the applicability to variant sequences, neglecting mutation-aware analysis that could enhance therapeutic relevance.

In this work, we integrate a wide array of computational tools on viral strain sequence, structure and property analyses to develop an antigen discovery pipeline with continuously expanding tool and data repositories for different viral families. The pipeline consists of three main stages. The first stage involves data collection and preprocessing, enabling the design of the antigen to be robust to sequence variation given viral strains under consideration. The second stage focuses on epitope prediction based on sequence and structure analyses. Followed by rigorous multi-step filtering and analyses, reflecting the central role of epitopes in triggering immune responses, the final stage designs and screens mutants at promising epitope regions as potential antigen candidates using existing functional score prediction tools, evaluating viability, immunogenicity, allergenicity, toxicity, and other relevant properties. To demonstrate its capability and effectiveness, we have tested our developed pipeline for antigen design across three different zoonotic viral families: coronavirus, phlebovirus, and alphavirus. Our preliminary results have shown promising potential in expediting future antigen development with broad efficacy in response to emerging pathogens.

## 2 The pipeline

In this section, we introduce the proposed antigen discovery pipeline in detail, as shown in Figure 1. The pipeline is initiated using a viral sequence and generates intermediate data, logs and the final outputs as it proceeds. The organization of these data components within the pipeline is illustrated in Figure 2.

**Figure 1.**
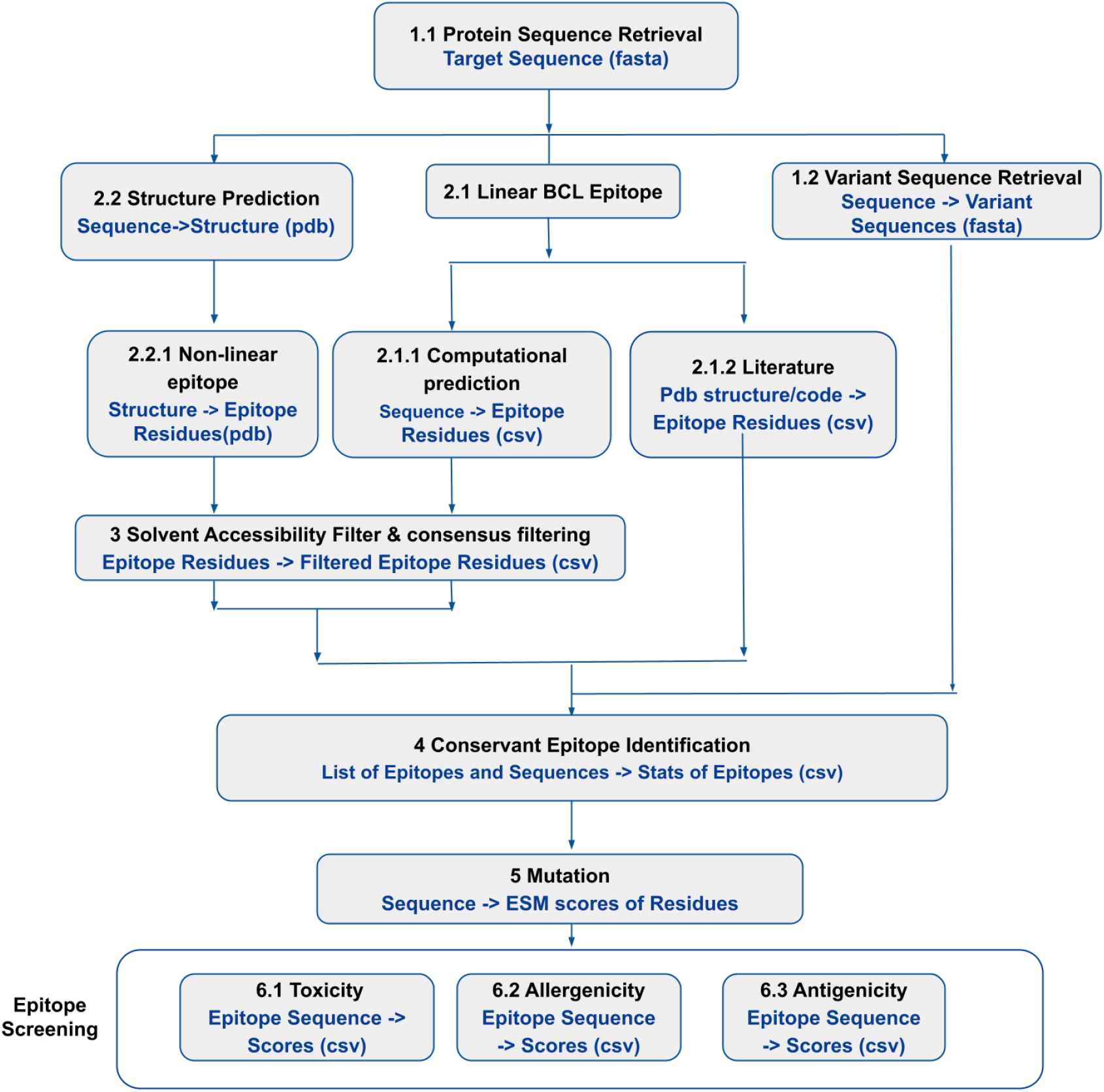
Illustration of the antigen prediction pipeline.

**Figure 2.**
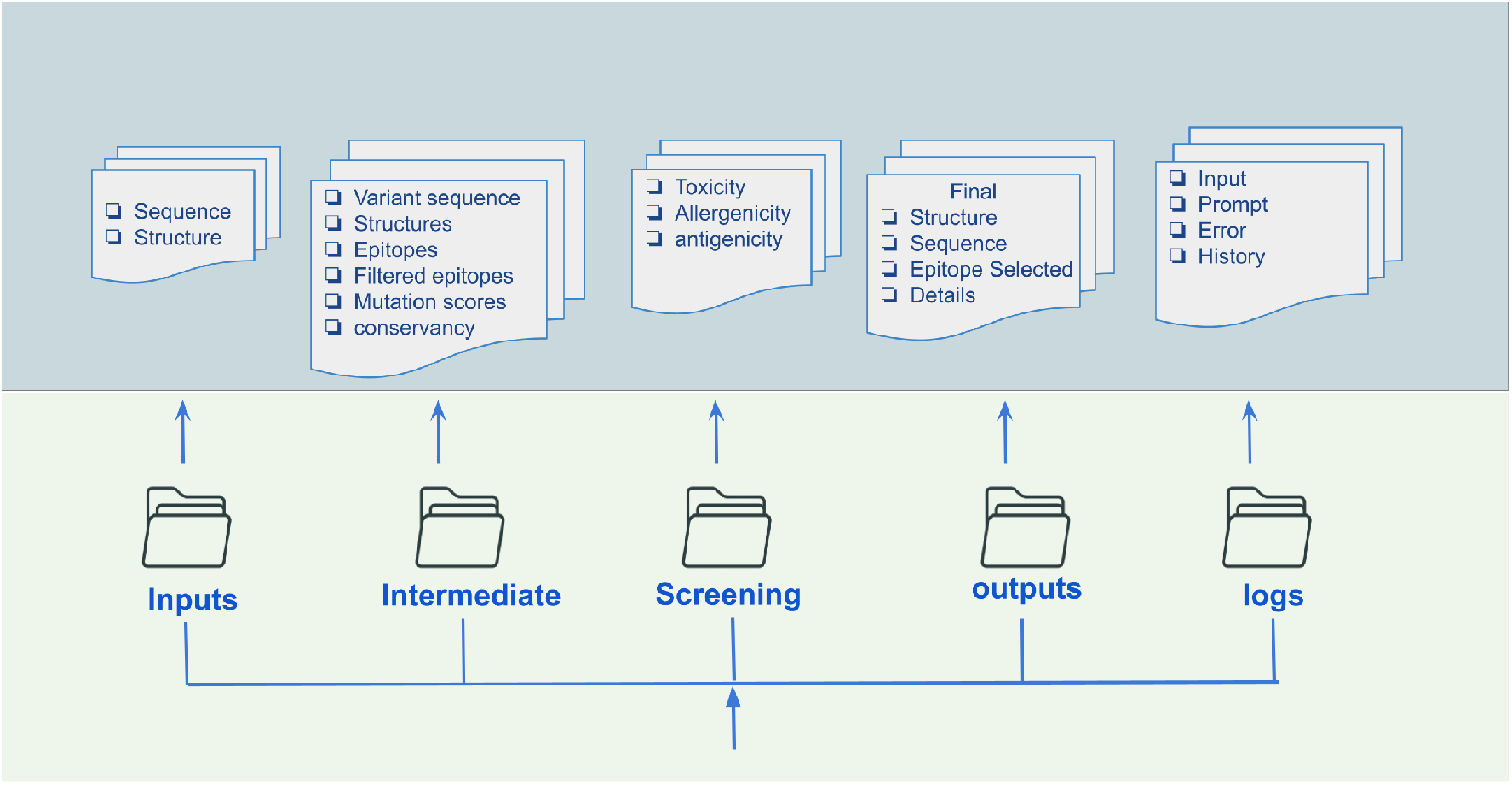
Data organization of the antigen prediction pipeline.

### 2.1 Data collection and processing

The data collection process involves both sequence searches and literature reviews. Since the ultimate goal of the pipeline is to design antigens with broad efficacy, it is essential to gather comprehensive sequence information from the target viral family. However, viral families differ significantly in their genetic variability, which influences downstream steps such as epitope prediction and mutation strategies. Therefore, the antigen design must incorporate the genetic variability of the target sequence.

### 2.2 Epitope analysis

In the current pipeline, we focus on B-cell epitopes, both linear and non-linear. Linear epitopes can be predicted using sequence-based epitope prediction tools, such as BepiPred3 [8], BepiBlast [40] and BepiPred2 [26]. On the other hand, we also predict protein structures using AlphaFold2 (AF2) [27] and AlphaFold3 (AF3) [1], if no experimental results are found in the PDB [4]. The structures are used to identify non-linear epitopes.

These predicted structures serves as inputs to DiscoTope2 [28] and DiscoTope3 [21], which output both predicted linear and non-linear epitopes. The output of these tools includes a binary classification that indicates if a residue is an epitope, along with its confidence score. In addition to sequence-based and structure-based epitope analysis, we can also resort to the literature to extract the structure of the virus of interest and its related structure from the Protein Data Bank (PDB).

The number of the predicted and literature-derived epitopes can be quite large. To reduce the number and increase the confidence of candidates for downstream analyses, we introduce a consensus approach. In this approach, we retain only those predicted epitope residues that are supported by multiple tools. For instance, the combined epitope classification (Comb) contains epitope residues that have been identified by at least one tool. In contrast, Consensus 2 (Cons2) and Consensus 3 (Cons3) require support from at least two or three tools, respectively. This strategy reduces reliance on individual tools and enhances the robustness of epitope selection.

### 2.3 Solvent Accessibility and Consensus Filtering

To further filter the candidate epitopes, we investigate the solvent accessible surface areas (SAS) using the combined results of two tools–FreeSASA and NetSurfP-3 [20]. NetSurfP-3 is a deep learning tool; it takes a sequence as input and provides residue-wise predictions of whether a given residue is exposed or not. In contrast, FreeSASA takes the structure of the target as input to calculate the exposed area. A threshold of 20 *Å*^*2*^ has been enforced to classify a residue as exposed. Traditionally, many thresholds have been used but in our pipeline, we have considered a stricter one.

Glycosylation sites are also excluded from the candidate epitopes as these sites have reduced accessibility [15] due to the binding of glycan molecules. These sites are identified using NetNGlyc - 1.0 [16]. After these filtering steps, the epitopes extracted from the literature are introduced to the pool of combined, consensus 2 and consensus 3 epitopes; the resulting pool of these epitopes are denoted as suggested lenient, suggested strict and suggested very strict respectively.

### 2.4 Mutation and Epitope Screening

The lack of reliable in silico tools forecasting viral evolution and predicting neutralization scores (e.g., IC50) limits our ability to directly optimize this antigen design computationally. Beyond neutralization, we aim to design antigens to be safe–non-toxic, non-allergenic and likely to fold correctly and be expressed properly. Computational predictors are available for these properties: Vaxijen 3.0 [13] for antigenicity, ToxinPred 3.0 [41] for toxicity, AlgPred 2 [45] for allergenicity, and the Evolutionary Scale Model (ESM) [18, 19, 31, 42] for predicting natural expression. Optimizing these objectives instead, we can reduce the search space and identify high-quality candidates, which are critical for success in non-convex optimization problems.

#### 2.4.1 Conservancy Analysis

The conservancy of the predicted epitopes is evaluated using the IEDB conservancy tool [5, 49], which has been integrated into the pipeline. The tool takes a list of variant sequences and a list of epitopes as input and outputs conservancy metrics. These include the percentage of protein sequence matches at identity *≤* 100%, as well as the minimum and maximum identity values of each epitope across the sequence pool. Epitope sequences shorter than three residues are discarded during this step. These findings are used to enhance the final selection of candidate epitopes.

#### 2.4.2 Mutation screening

To enhance the therapeutic properties of the predicted epitopes, mutation strategies were applied using a pretrained ESM model [35].

ESM [31] is trained as a masked language model, so it enables us to compute the probability ratio between mutated residue and *wildtype* residue, treating the other residues as *contexts*. Mathematically, the ESM score is defined as follows:

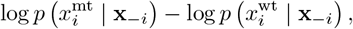

where *x*_*i*_ denotes the residue at position *i*, **x** denotes a sequence of residues, and **x**_*−i*_ means the sequence whose *i*-th position is masked.

We first introduce mutations on the epitope residues, with the goal to boost the properties mentioned above, so that the antigenicity score would become better, except for if it is not 100% immunogenic as per the VaxiJen3.0 tool, which we will refer to as the desired antigenicity. Simultaneously, the introduced mutations should not make the epitope more toxic or allergenic. To achieve this, we use multiple threshold values, for instance, 85th, 90th, 95th, and 99th percentile values of the ESM-scores to mutate the candidate epitopes. In this phase, the focus is narrowed down to a specific domain based on the literature.

## 3 Results

In this section, we showcase the efficacy of our developed antigen design pipeline for three viruses: SARS-CoV-2 in coronavirus, RVFV in phlebovirus, and MAYV in alphavirus. We first validate the pipeline with SARS-CoV-2, and provide case study results on RVFV and MAYV.

### 3.1 Detection of Conserved Epitopes in SARS-CoV-2

Due to the availability of vast experimental data, SARS-CoV-2 was chosen to validate the pipeline. We took Wuhan-Hu-1 (YP 009724390), obtained from GenBank, as our starting target sequence for antigen design, and the corresponding structural were retrieved from the Protein Data Bank (PDB) [6]. Using the target sequence, we queried and collected 100 related sequences. After removing entries with missing residues, 88 complete sequences remained for the analysis.

To validate the pipeline, epitopes were extracted from the antigen-antibody complex structures available in the PDB. These literature-derived epitopes were then mapped to the Wuhan-Hu-1 sequence with the help of sequence alignment. We define epitope–paratope pairs as residue pairs in the antigen-antibody complex whose respective alpha carbon atoms (C*α*) are within 8Å of each other. This distance-based criterion is commonly used in protein–protein interaction studies to identify interface residues. [44, 53, 54].

During the epitope analysis stage, epitopes were predicted using both the sequence and structural information. Notably, no literature epitopes were added to the pool of epitope candidates. This resulted in 1169 candidate eptiope residues in total.

After applying SAS filtering, the number of candidate epitopes decreased across all consensus levels: from 1169 to 1028 (−12.06%) for the Comb set, from 483 to 440 (−8.9%) for Cons2, and from 149 to 137 (−8.05%) for Cons3. Applying a consensus threshold of 2 reduced the SAS-filtered set from 1,028 to 440 residues (−57.19%), and applying a consensus of 3 further reduced this to 137 (−68.86%). Furthermore, we excluded glycosylation sites, which led to the reductions in candidate epitopes: from 1028 to 953 (*™*7.30%) for the combined set (Comb), from 440 to 400 (−9.09%) for Cons2, and from 137 to 127 (−7.3%) for Cons3, respectively. For the rest of the experiment, we focused on the candidates from Cons3 to decrease false positives.

We further restricted the search region to the Receptor-Binding Domain (RBD) of the spike protein. Empirically, it is known to be a crucial binding site of human antibodies [39]. This further reduced the candidate epitopes to 68 from 127 (−46.45%) by Cons3 filtering (Figure 3B). A subsequent conservancy analysis identified 41 conservant epitope residues in the RBD (Figure 3C). We extracted 36 validated epitope residues form the literature [29, 43, 51], of which 16 are present in our candidate pool and 11 of them are in the conserved regions (Figure 3D). More details about conserved epitopes are presented in Table 2.

**Figure 3.**
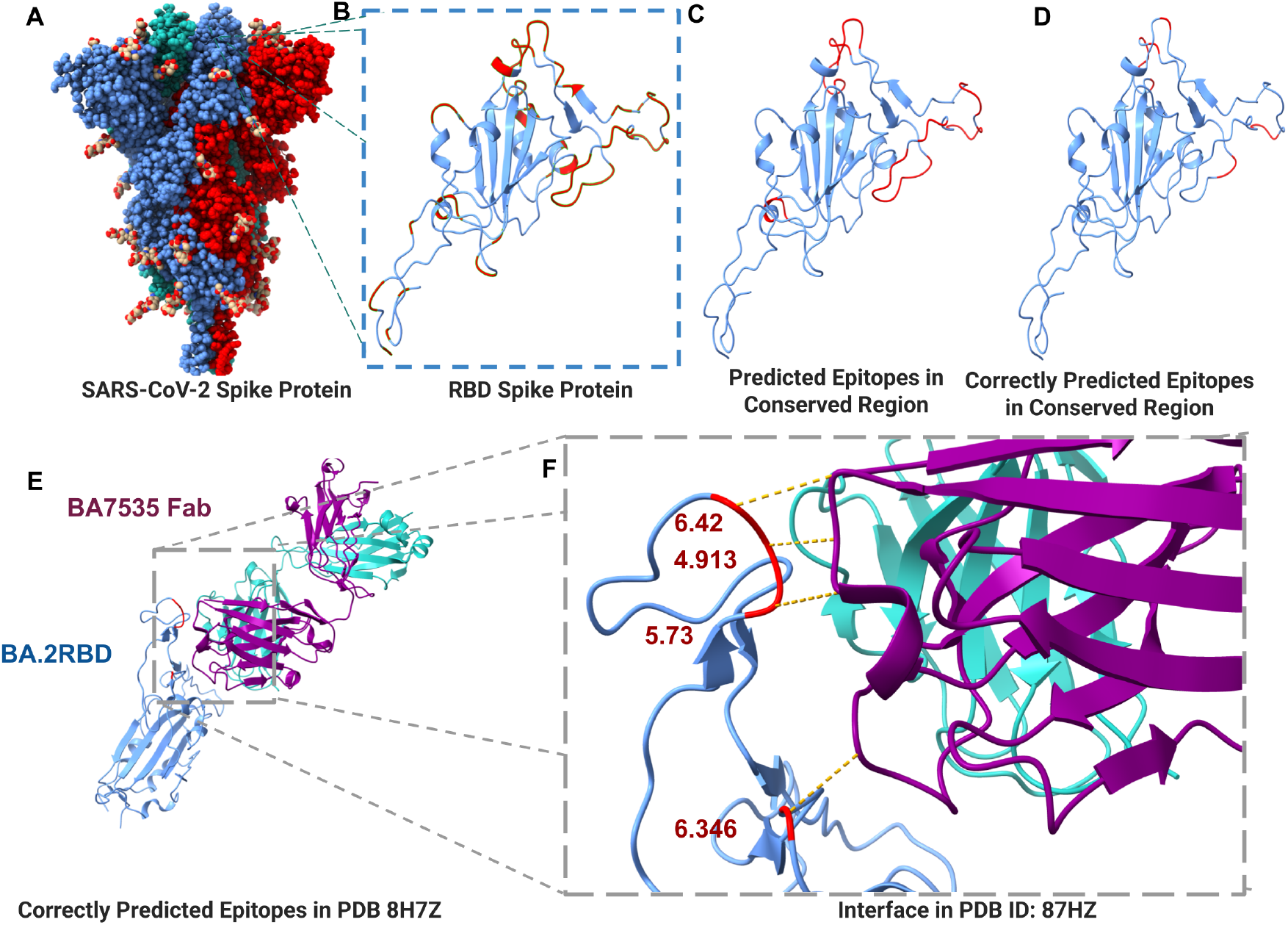
Epitope Analysis in SARS-COV-2. A) Structure of SARS-CoV-2 spike protein (Wuhan-Hu-1); B) Focused structure visualization of the RBD region of spike protein colored in sky blue and all the predicted epitopes in cons3 highlighted in red; C) Predicted epitopes in conserved region highlighted in red; D) Predicted conserved epitopes mapped (highlighted in red) that overlap with known epitopes, which were used as ground truth for validation; E) The predicted conserved epitopes (highlighted in red) that overlap with known epitopes in PDB: 8H7Z. BA755 Fab is colored in purple and cyan; F) Focused binding interface of BA7535 Fab and BA.2RDB. Some of the interactions of residues are shown (in yellow) and their interchain distance (Ca-Ca) in Å

**Table 1:** Epitope counts before and after SAS filtering, with suggested thresholds.

| 2*Virus | Before SAS |  |  | After SAS |  |  | Lit. | Glyco | Suggested |  |  |
| --- | --- | --- | --- | --- | --- | --- | --- | --- | --- | --- | --- |
|  | Comb | Cons2 | Cons3 | Comb | Cons2 | Cons3 |  |  | Lenient | Strict | Very strict |
| **SARS-CoV-2** | 1169 | 483 | 149 | 1028 | 440 | 137 | 0 | Yes | 953 | 400 | 127 |
| **RVFV** | 804 | 381 | 163 | 725 | 373 | 163 | 82 | No | 735 | 398 | 217 |
| **MAYV** | 233 | 109 | 34 | 208 | 101 | 34 | 20 | No | 222 | 117 | 54 |

**Table 2:**
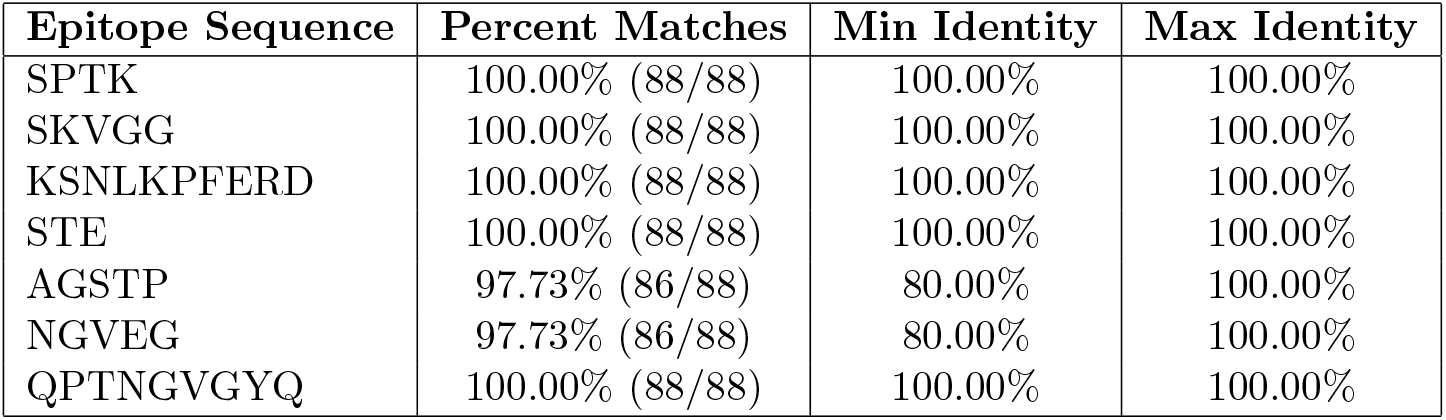
Conservation of selected epitope sequences across related protein sequences of the RBD region SARS-CoV–2.

Importantly, we found that the epitopes suggested by the pipeline overlapped with highly conserved regions targeted by the neutralizing antibody BA7535 (PDB ID: 87HZ) [51]. BA7535 has been empirically validated that to exhibit high neutralizing potency across a broad spectrum of SARS-CoV-2 variants including Alpha, Beta, Gamma, Delta, Omicron BA.1-BA.5, BF.7, CH.1.1, XBB.1, XBB.1.5, XBB.1.9.1, and EG.5 strains. Figures 3E & F visualize the binding interface of BA7535 and SARS-CoV-2 BA.2 RBD. A total of 10 epitope residues were found from the validated structure. Of the 10 epitope residues, six are present in our candidate pool, and four of these lie within conserved regions.

Finally, the pipeline proceeded with the mutation design the and epitope screening. Table 3 summarizes the number of candidate epitopes retrieved at each ESM score percentile, before and after filtering screening. As a result of rigorous screening procedure only 10 candidates are left from 201 mutated epitope pool for the 90th perecentile value of ESM score. This demonstrates the capability of the pipeline to reduce the search space.

**Table 3:** Number of epitopes with the desired properties at various percentile thresholds of ESM scores for RVFV and MAYV.

| Percentile threshold of ESM score | Before screening |  |  | After screening |  |  |
| --- | --- | --- | --- | --- | --- | --- |
|  | SARS-CoV-2 | RVFV | MAYV | SARS-CoV-2 | RVFV | MAYV |
| 85th | 354 | 619 | 423 | 13 | 16 | 27 |
| 90th | 201 | 603 | 252 | 10 | 16 | 16 |
| 95th | 62 | 345 | 112 | 6 | 9 | 8 |
| 99th | 6 | 73 | 19 | 1 | 2 | 1 |

Our results demonstrate that our pipeline can effectively search the antigen search space to identify the neutralizing epitope residues of SARS-CoV-2. It can also highlight the conserved regions across multiple strains of a virus family for broader effective antigen design. Moreover, it suggests mutations that may enhance the immunogenic property of the antigen. Collectively, these findings underscore the potential of the pipeline for real-world applications in antigen design and immuno-therapeutic development.

### 3.2 Case studies on RVFV & MAYV

We further test our antigen design pipeline for RVFV and MAYV. The target sequence for each virus was obtained from GenBank, accession code of ABD38813.1 for RVFV and AF237947.1 for MAYV, respectively.

Given the corresponding target sequence, for MAYV, we have retrieved 66 similar viral strain sequences; and after removing the sequences with missing residues, 59 remained. For RVFV, 375 sequences were retrieved, and after removing the sequences with missing residues, 351 were left. Multiple sequence alignment (MSA) was performed by CLUSTALW [47] for each set of the virus sequences. These MSAs were then used in DiMA [46]. to measure sequence diversities with default settings. Figure 4 displays the sequence diversity analysis results. The goal of the analysis was to be aware of the sequence diversity and also to focus on more conserved regions with lower diversity.

**Figure 4.**
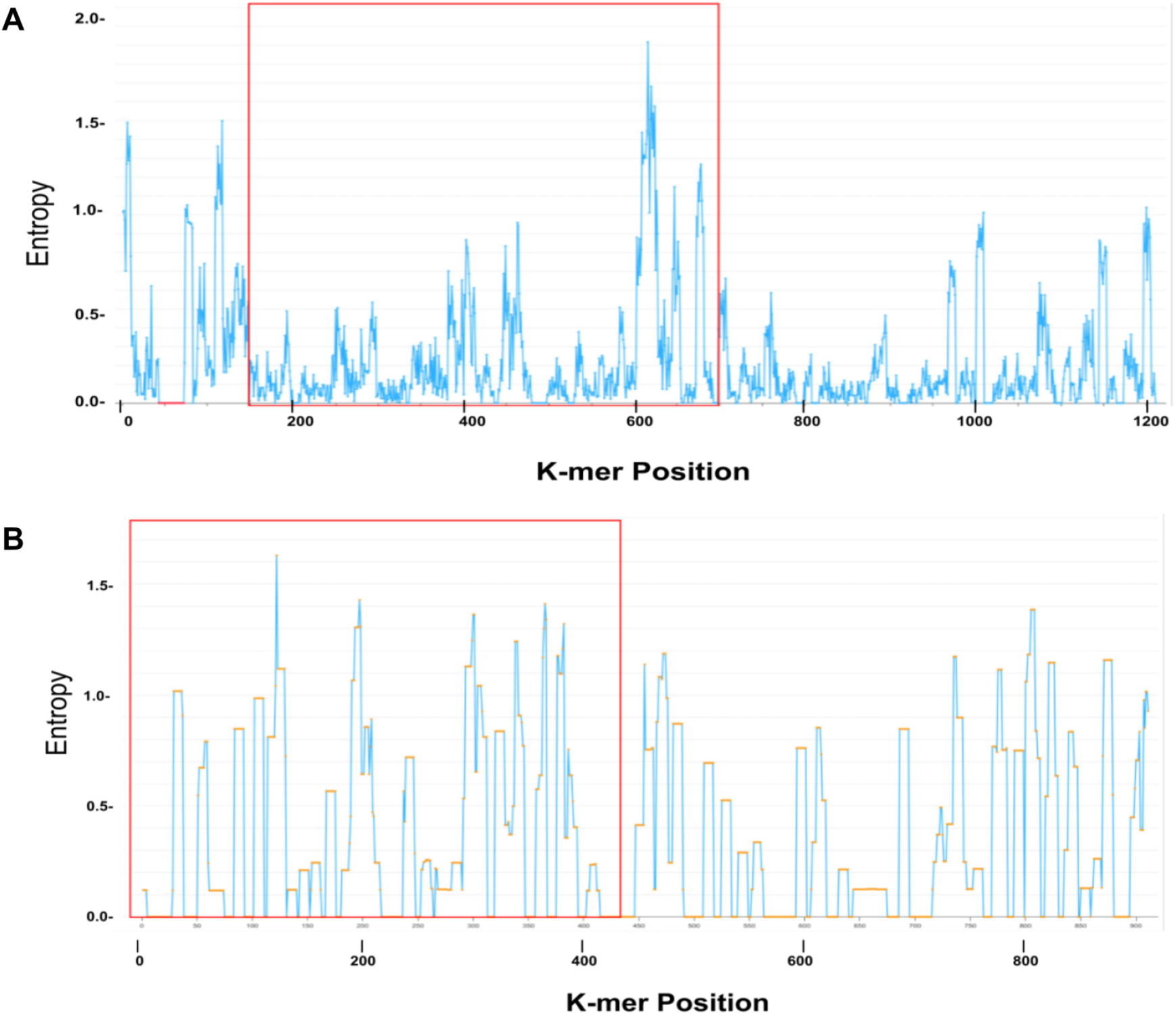
Sequence diversity analysis using DiMA with the resulting sequence diversity plots of A) RVFV and B) MAYV, respectively. The y-axis is the entropy, which is directly proportional to sequence diversity, and the x-axis is K-mer Position, which is analogous to residue index. In A), the red box represents the Gn-region of the M-segment of the RVFV, which has been focused as antigen; and in B) the red box represents the E2-Glycoprotein of the MAYV.

**Figure 5.**
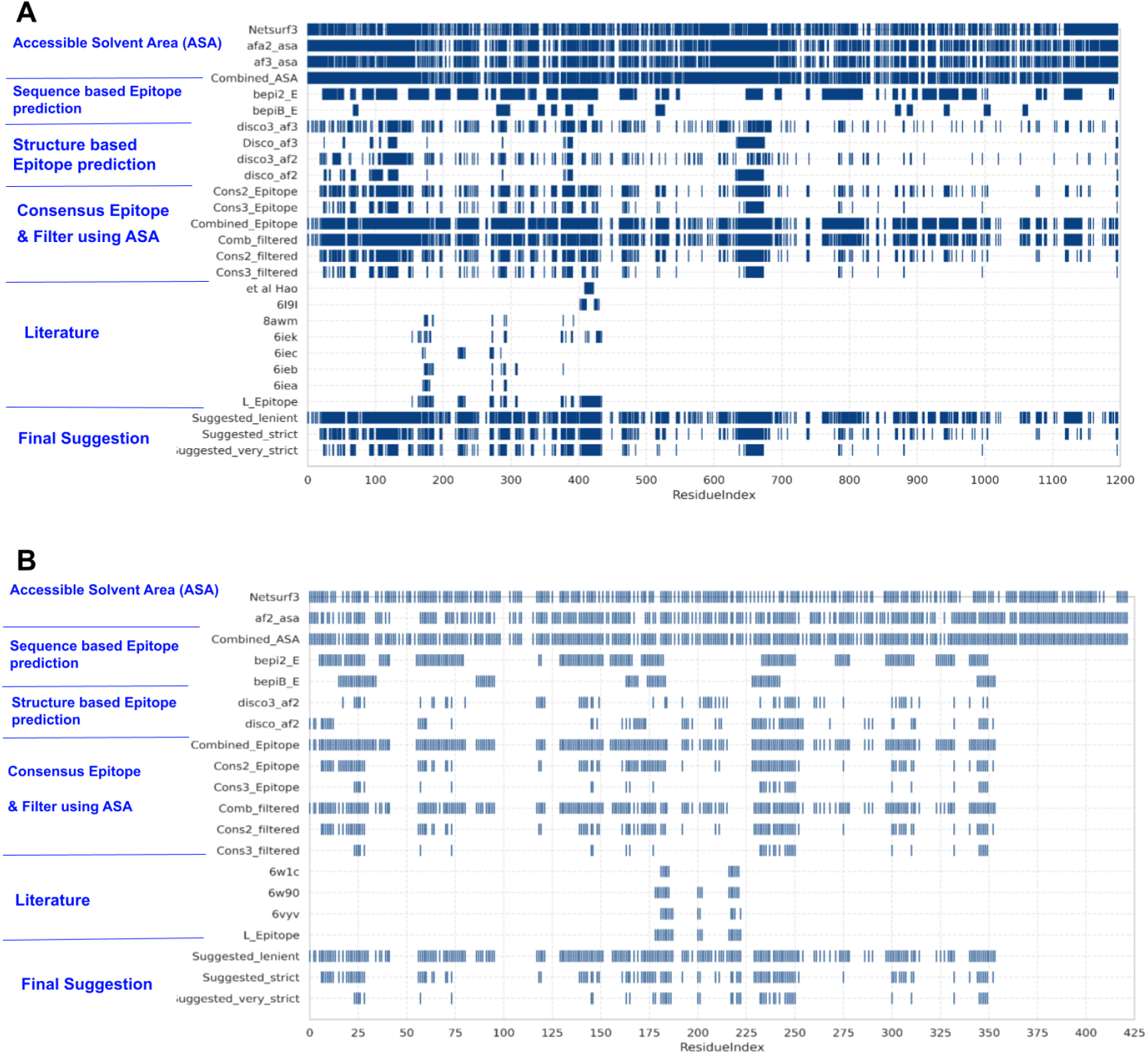
Heatmap of the epitope candidate filtering: A) RVFV, B) MAYV.

**Figure 6.**
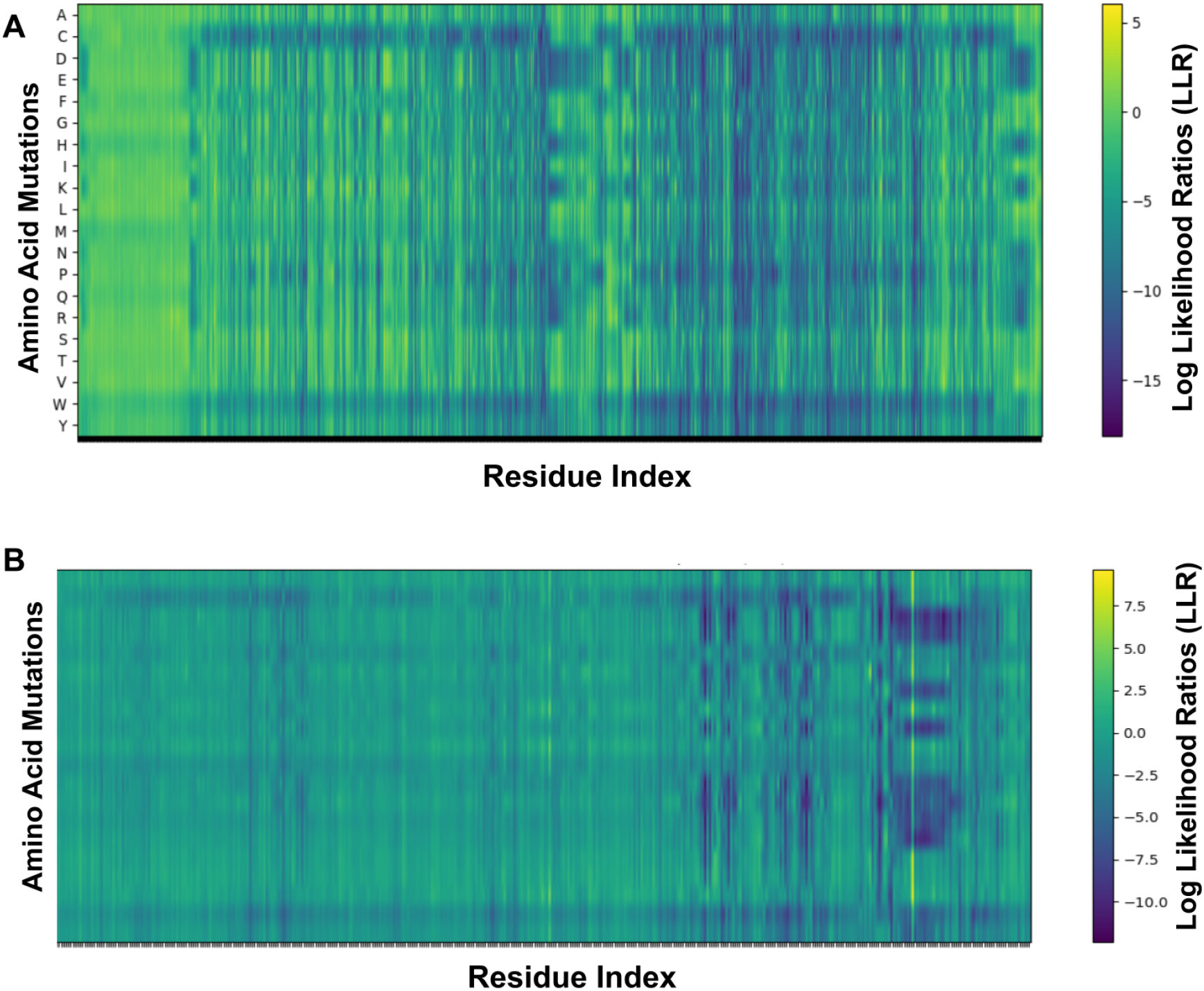
Heatmaps of the ESM scores for every single mutation: A) RVFV, B) MAYV.

The next step is the B-cell epitope prediction using multiple tools. It was noteworhty that for RVFV, we used structures predicted by both AlphaFold2 (AF2) and AlphaFold3 (AF3). In contrast, for MAYV only the predicted structure using AF2 was used, as the prediction was very similar to AF3. Consequently, the sequence and the structure was used to predict the epitopes for each virus. As a result the total number of predicted epitope residues were 233 for MAYV and 804 for RVFV.

Table 1 summarizes the progressive refinement of candidate B-cell epitope residues leveraging a multi-step filtering strategy. It presents the effect of applying filters such as SAS, varying the consensus threshold, and the final suggestions. The initial set of epitopes, presented under “Before filter using SAS,” listed under all the epitope residues predicted by multiple methods.

To refine the pool of candidates, the buried residues were removed using the SAS prediction. This strategy reduced the “Comb” candidate number of epitope residues by 10.73% for MAYV and 9.83% for RVFV, respectively. However, the reduction rate decreased as the degree of consensus was increased, suggesting that a higher degree of consensus consists of a more refined pool of candidates. Notably, the consensus-based approach had a more significant impact on the candidate pool. After applying the Cons2 threshold, the number of candidates decreased by 51.4% for MAYV and 48.55% for RVFV, respectively, even after applying SAS. Applying the Cons3 threshold to the pool of Cons2, it decreased by 66.3% for MAYV and 55.1% for RVFV, respectively. Comprehensive details about the scores are available in the appendix. In addtion, The validated candidates extracted from the literature [2, 7, 17, 38, 50] were combined into the pool, which enhances the candidate pool quality. Finally, we are left with three different classes of suggested epitopes.

For the strict suggestion, the number of initial candidates decreased from 233 to 117 (−49.80%) for MAYV and from 804 to 398 (−50.50%) for RVFV. This demonstrated the capability of the pipeline to reduce the large space of candidates to a manageable number that can be adjusted based on the user’s needs. For instance, for the next part of the experiments, we used the epitope candidates using the strict approach since BepiPred3 was not used for this analysis.

Furthermore, mutation analysis was performed, and at multiple thresholds, the number of candidate epitope residues before and after screening is summarized in Table 3. From here, the focus of search was shifted to the E2-glycoprotein of MAYV and the glycoprotein-n (GN) region of RVFV, respectively. Following a rigorous screening process, only a small subset of candidate epitopes was left. From these filtered pools of candidates, conservation analysis can be incorporated to focus on the conserved epitope regions.

We observed a particular promising epitope candidate in RVFV at the residue index 563 of the target sequence, which exhibited several desirable properties. This epitope is neighboring to a residue that overlaps with one of the mutations (index 566) identified in the recombinant MP-12 strain (MP-12) sequence, GenBank accession code DQ380208. It is a vaccine strain of the RVFV, developed specifically for vaccination purposes [23, 33]. Further analysis suggested that the ESM score of that specific mutation was the top-4 value out of all the ESM scores in the RVFV wild-type m-segment sequence. This provides crucial insight into using ESM values as a guide for mutation design.

## 4 Discussion

This pipeline employs hierarchical filtering to reduce the search space and uses mutation screening to boost desirable properties. In addition, it focuses on epitopes located in conserved regions of target-related sequences. Moreover, it introduces functionalities absent in related works such as mutation design, multi-step filtering and conservation-aware prioritization. The pipeline exhibited promising results when applied to the SARS-CoV-2 and demonstrated that ESM can be used reliably to guide mutation.

Altogether, the pipeline has the potential to support the development of antigens capable of eliciting a broad and effective immune response. While promising, it is important to note that the pipeline is in an early stage of development. It relies on third-party tools, some of which need to be improved due to limited effectiveness. Another key limitation is the candidate screening process, which could be improved through exploring multi-objective optimization approaches that integrate multiple scoring metrics. Also, the final candidates of the pipeline are not validated experimentally. The integration of ground-truth data such as serum assay results into the pipeline could have been used to boost its performance.

Despite the limitation, this provides a solid starting point for an open-source antigen design pipeline. Furthermore, it can be continuously modified as the field evolves. Incorporating agentic system can increase the pipeline’s flexibility, enabling it to dynamically tailor the selection of tools used in the pipeline based on the user’s needs.

## 5 Conclusion

In this work, we have presented a robust integrated computational pipeline for designing antigens that can address the evolving viral diseases. Our evaluations of the pipeline on SARS-CoV-2 and RVFV have shown promising performance. For example, the pipeline has predicted epitope residue mutations in an experimentally validated epitope of SARS-CoV-2. When designing antigens for RVFV, one of the predicted potential epitopes overlaps with one of the introduced mutations in the previously developed MP-12 strain, a promising vaccine candidate reported in literature [24]. Experimental validation of the designed antigens for the desired neutralizing responses of RVFV and MAYV can provide stronger evidence of the effectiveness of our designed pipeline and constituent tools.

While promising, our current pipeline has a few limitations, especially on the generalizablity and efficacy of the current property prediction and mutation engineering tools, which shall be improved in future research for better mutation-level sensitivity and explainability. Future research and more experimental data are also valuable to assess the pipeline for broad efficacy of epitope analysis and antigen design given different viral families. Recent agentic workflow development can also improve user interactions to tailor a pipeline as per the user’s need. Ultimately, with continued refinement and validation, this pipeline offers a promising foundation for open-source, data-driven antigen design.

## 7 Author contributions statement

R.S.R., B.J.Y. and X.Q. conceived the experiment(s), R.S.R. conducted the experiment(s). R.S.R., B.J.Y. and X.Q. analyzed the results. R.S.R., B.J.Y., and X.Q. wrote the manuscript. T.I., S.W., N.V. provided guidance and feedback regarding the MAYV and RVFV antigen design. All authors contributed to the discussions, reviewed and edited the manuscript.

## 8 Acknowledgments

This work has been supported by the ARPA-H Award #1AY1AX000053.

